# Shining a light on Laurentian Great Lakes cisco (*Coregonus artedi*): how ice coverage may impact embryonic development

**DOI:** 10.1101/2021.03.23.436622

**Authors:** Taylor R. Stewart, Mark R. Vinson, Jason D. Stockwell

## Abstract

Changes in winter conditions, such as decreased ice coverage and duration, have been observed in the Laurentian Great Lakes for more than 20 years. Such changes have been hypothesized to be linked to low *Coregonus* spp. survival to age-1 as most cisco (*Coregonus artedi*) populations are autumn spawners whose embryos incubate under ice throughout the winter. The quantity of light during winter is regulated by ice coverage, and light affects embryo survival and development in some teleosts. We experimentally evaluated how cisco embryos from lakes Superior and Ontario respond to three light treatments that represented day-light intensity under 0-10, 40-60, and 90-100% ice coverage. Embryonic response measures included two developmental factors (embryo survival and incubation period) and two morphological traits (length-at-hatch and yolk-sac volume). Embryo survival was highest at the medium light treatment and decreased at high and low treatments for both populations, suggesting cisco may be adapted to withstand some light exposure from inter-annual variability in ice coverage. Light intensity had no overall effect on length of incubation. Increasing light intensity decreased length-at-hatch in Lake Superior but had no effect in Lake Ontario. Yolk-sac volume was positively correlated with increasing light in Lake Superior and negatively correlated in Lake Ontario. Contrasting responses in embryo development between lakes suggests differences in populations’ response to light is flexible. Our results provide a step towards better understanding the high variability observed in coregonine recruitment and may help predict what the future of this species may look like under current climate trends.

## Introduction

Freshwater whitefishes, Salmonidae Coregoninae (hereafter coregonines) play important economic (Ebener et al., 2008) and ecological (Lynch et al., 2010; Nyberg et al., 2001; Stockwell et al., 2014) roles throughout the northern hemisphere, but populations have declined over the past century (Eshenroder et al., 2016). Historical coregonine declines were attributed to overfishing, invasive species, habitat alterations, and competition (Anneville et al., 2009; Rosinski et al., 2020; Stockwell et al., 2009). More recently, coregonine populations worldwide have experienced declines due to highly variable recruitment and low survival to age-1 (Lepak et al., 2017; Nyberg et al., 2001; Parks and Rypel, 2018) which have been associated with climate-induced changes in early-life stage environments (Nyberg et al., 2001). However, an underlying mechanism between changing lake environments and coregonine year-class strength has yet to be established.

Year-class strength in most fish species, including coregonines, is thought to be established prior to the end of the first season of growth (Cushing, 1990; Hjort, 1914; Karjalainen et al., 2015). Most coregonines are autumn spawners whose embryos incubate under ice throughout the winter (Karjalainen et al., 2000; Stockwell et al., 2009). Embryos are static, which leaves them vulnerable to predation (Stockwell et al., 2014) and unable to evade detrimental changes in winter environmental conditions (Pepin, 1991). Changes in winter conditions, such as decreased ice coverage and duration, that have been observed over the past 20+ years (Austin and Colman, 2007; O’Reilly et al., 2015; Sharma et al., 2019), could alter developmental rates, embryo survival, and time of hatching (Karjalainen et al., 2015). Potential mechanisms by which ice coverage influences coregonine embryonic development include reduced physical wave action (Austin and Colman, 2007; Nguyen et al., 2017; Walter et al., 2006; Wang et al., 2010), stabilized winter and spring water temperatures (Magnuson et al., 1997; Winslow et al., 2017), and the amount of sunlight reaching the lake bottom (Bolsenga and Vanderploeg, 1992; Hampton et al., 2015).

Photoperiod is the most consistent abiotic factor in nature (Ruchin, 2020) and can regulate fish development phenology, behavior, and physiology (Ruchin, 2007; Villamizar et al., 2011). The length of photoperiods characterize circadian rhythms and ensure that biological processes are synchronized with the environment (Gaston et al., 2013; Marchesan et al., 2005; Ruchin, 2020). In seasonally ice-covered lakes, winter lake light levels are regulated by ice coverage and snow depth (Bolsenga and Vanderploeg, 1992; Hampton et al., 2015). Ice can reduce light transmittance to 62% under clear ice, and to ≤ 10% under snow covered ice (Bolsenga and Vanderploeg, 1992).

Salmonid embryos incubated under elevated light levels had higher mortality and deformity rates, slower formation of cartilaginous skeletal elements, decreased time to hatching, smaller size-at-age, and accelerated development after organogenesis (Chernyaev, 2007; Eisler, 1961, 1958; Kwain, 1975; MacCrimmon and Kwain, 1969). However, other teleosts (e.g., turbot *Scophthalmus maximus*, Atlantic halibut *Hippoglossus hippoglossus*, brown-marbled grouper *Epinephelus fuscoguttatus*) exhibit opposite responses, or no response, to manipulated light illumination during incubation (Iglesias et al., 1995; Mangor□Jensen and Waiwood, 1995; Ruchin, 2020; Seth et al., 2014). To our knowledge, no previous work has examined the effects of light on coregonine embryos from North America.

We experimentally evaluated how cisco (*Coregonus artedi*) embryos responded to different photoperiod intensities, as a proxy for different ice coverages. We hypothesized that exposure to elevated light intensity (a proxy for low ice coverage) decreases embryo survival and accelerates embryogenesis, resulting in earlier hatching, larger yolk-sac volume, and shorter length-at-hatch. Our objective was to identify the extent to which light influences cisco embryo survival, incubation duration, and length and yolk-sac volume at hatching. We expected populations adapted to lower light levels (high ice coverage) would experience a greater magnitude of change as light intensity increases.

## Methods

### Ethics

All work described here was approved for ethical animal care under University of Vermont’s Institutional Animal Care and Use Committee (Protocol # PROTO202000021).

### Study Species and Locations

Mature cisco were collected from the Apostle Islands, Lake Superior (46.85°, -90.55°) and Chaumont Bay, Lake Ontario (44.05°, -76.20°) in December 2019. Lake Superior cisco were collected at an open lake location at depths between 15 and 50 m. Lake Ontario cisco were collected in a shallow, protected bay on rocky shoals at depths between 2 to 5 m. Egg deposition has been confirmed in Chaumont Bay (George et al., 2017; Paufve et al., 2020). No direct evidence of spawning has been observed in Lake Superior and thus we are using the presence of ripe adults at our collection location as a proxy for a spawning location. We acknowledge that spawning and the embryo incubation location could be different, but previous literature suggests that spawning in Lake Superior occurs at depths of 30-200 m (Dryer and Beil, 1964; Eshenroder et al., 2016; Stockwell et al., 2009). Historical (1973-2020) ice conditions over the sampled spawning locations varied between lakes with the shallower, more protected Lake Ontario spawning site having more consistent ice coverage between January and March than the deeper, open location in Lake Superior (Figure 1). The different spawning habitats provide a contrast in light levels that coregonine embryos from each population would naturally experience because maximum light availability decreases with depth (Fleming-Lehtinen and Laamanen, 2012; Preisendorfer, 1986; Ramus et al., 1976; Secchi, 1864) and winter light availability is further restricted by ice and snow conditions (Bolsenga and Vanderploeg, 1992; Hampton et al., 2015).

**Figure 1.**
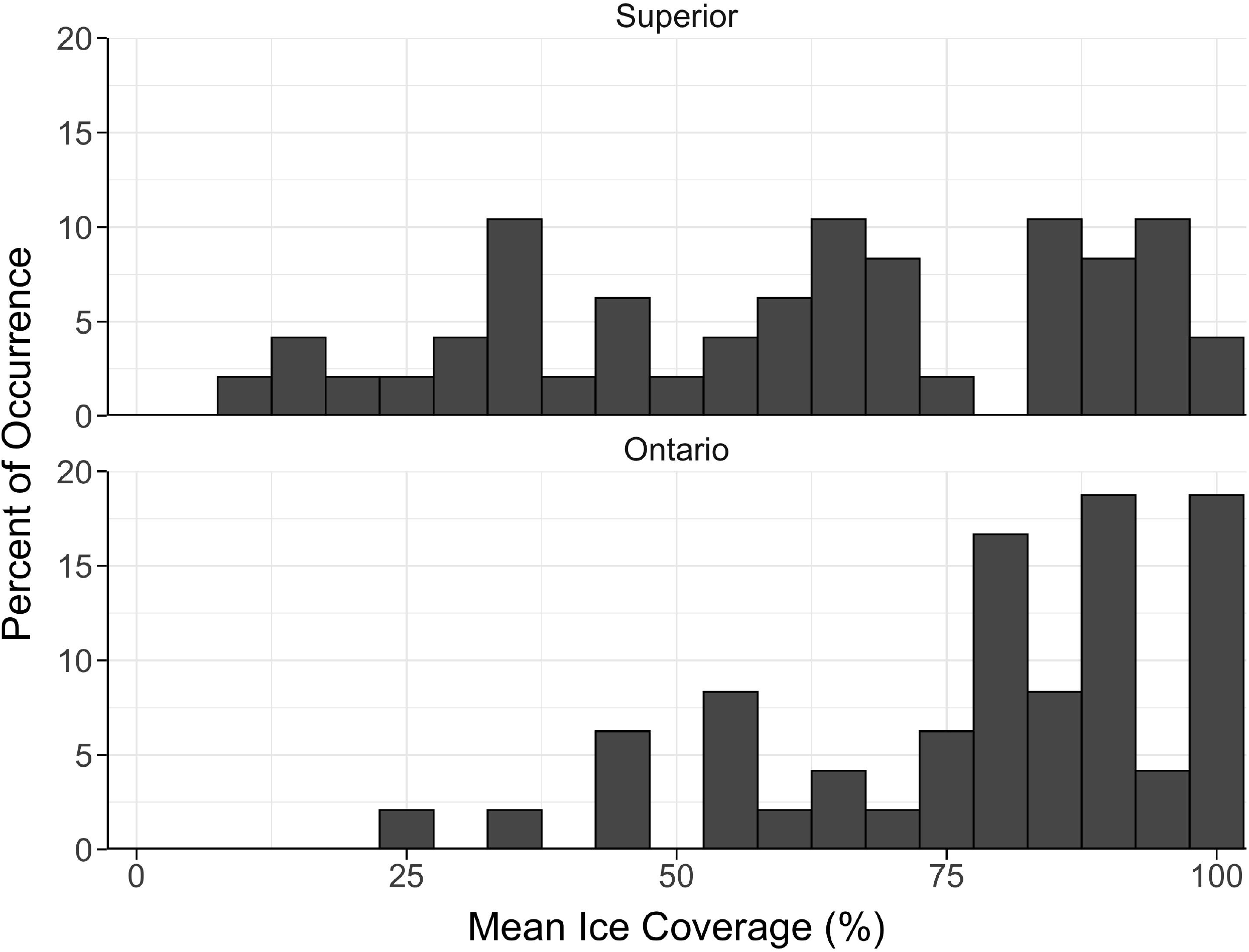
Histogram of annual mean ice coverage between 1-Jan and 15-Mar from 1973-2020 for the sampling location in Lake Superior (top) and Lake Ontario (bottom). Ice coverage data was obtained from the U.S. National Ice Center (usicecenter.gov/).

### Crossing Design and Fertilization

The design is fully described in Stewart et al. (2021). Briefly, gametes were stripped from 12 females and 16 males from each lake and artificially fertilized to create 48 families from each lake. Reconstituted freshwater medium (ISO 6341, 2012) was used during fertilizations and rearing to standardize the chemical properties of the water used between lakes. Embryos were transported to the University of Vermont in coolers by overnight shipping for Lake Superior samples and driven the same day for Lake Ontario samples. A temperature logger recorded air temperature inside the transport cooler (Lake Superior: mean (SD) = 2.80°C (0.21); Lake Ontario: mean (SD) = 3.28°C (0.37)). Total length, mass, and egg diameter were collected from the spawned adults. Fertilization success was determined by assessing 10 haphazardly selected embryos under microscopy (Oberlercher and Wanzenböck, 2016). If fertilization was low (<30%), the family was removed from the experiment.

### Rearing Conditions

Embryos were individually distributed into 24-well cell culture microplates and incubated in 2 ml of reconstituted freshwater (Stewart et al., 2021). A total of 36 embryos were used for each Lake Ontario and Lake Superior cisco family. Families were randomly distributed across three microplates (i.e., 12 eggs per family per microplate resulting in two families per 24-well microplate).

Microplates from each population were incubated under three light treatments (0.6-6.2 μmol m^-2^ s^-1^) that represented day-light intensity under 0-10 (low), 40-60 (medium), and 90-100 % (high) ice coverage (Table 1) and followed mean weekly photoperiods with gradual sunrise and sunset transitions. Light intensities for each treatment were chosen to mimic *in situ* winter, lakebed light measurements that were previously recorded with a photometer (JFE Advantech Co., Ltd. DEFI2-L) from Lake Superior (46.97°, -90.99°) at 10 m of water in 2016-17. No light intensity measurements were taken from Lake Ontario. Remote-sensing ice data (U.S. National Ice Center; usicecenter.gov) were used to quantify the daily percentage of ice coverage above the light sensor (Figure 2). Embryos were incubated at a constant target water temperature of 4.0°C in a climate-controlled chamber (Conviron^®^ E8; Table 2). Forced airflow was used in the climate-controlled chamber to ensure equal air circulation around the microplates and opaque, plastic sheeting was used to separate light treatments. Microplates were covered with transparent lids to minimize evaporation and rotated (i.e., orientation and position within the incubator) weekly. Water temperature and light intensity were recorded hourly with loggers (HOBO^®^ Water Temperature Pro v2 and JFE Advantech Co., Ltd. DEFI2-L) and daily mean values calculated (Table 1). During the hatch period, microplates were checked on a three-day cycle for newly hatched embryos. All hatched embryos were photographed ventrally (Nikon^®^ D5600 and Nikon^®^ AF-S DX 18-55mm lens) and then immediately preserved in 95% ethanol. Egg size at fertilization, total length-at-hatch, and post-hatching yolk-sac axes were measured from photographed images using Olympus^®^ LCmicro.

**Table 1.**
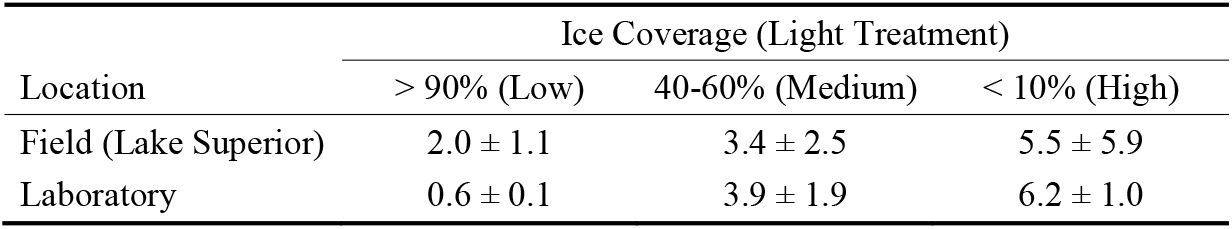
Mean daily ± SD light intensity (μmol m^-2^ s^-1^) for three ice coverage classes measured from Lake Superior and corresponding laboratory experimental light conditions used for both Lake Superior and Lake Ontario.

**Table 2.**
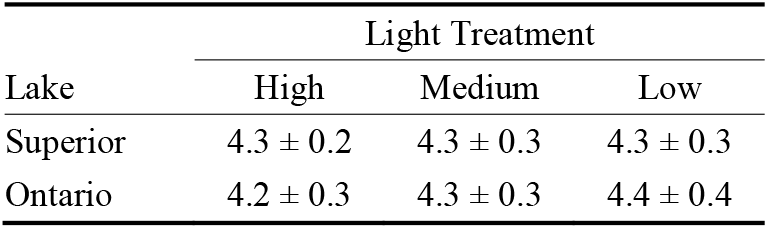
Mean daily ± SD water temperatures (°C) during embryo incubations from each light treatment for Lakes Superior and Ontario.

**Figure 2.**
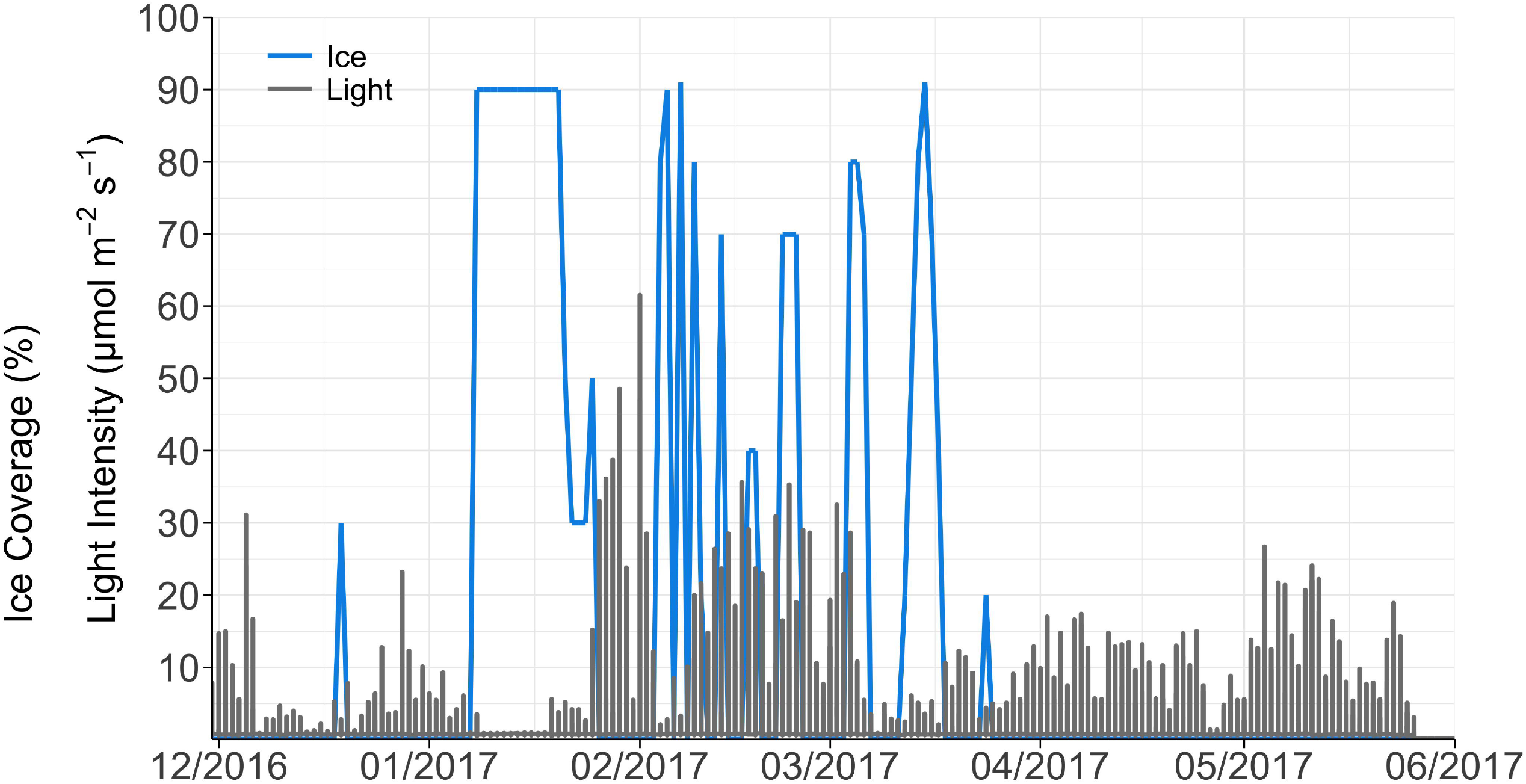
Daily ice coverage (%; blue line) and light intensity (μmol m^-2^ s^-1^; gray line) based on light sensors set at 10 m depth off Sand Island, Lake Superior. Ice coverage data above the sensor was obtained from the U.S. National Ice Center (usicecenter.gov/).

### Developmental and Morphological Traits

Embryo survival was estimated as the percent of embryos surviving between eye-up and post-hatch stages. Incubation period was assessed with two variables: the number of days from fertilization to hatching (days post-fertilization; DPF) and the sum of the degree-days to hatching (accumulated degree-days; ADD; °C). Total length-at-hatch (LAH; mm) and yolk-sac volume (YSV; mm^3^) were measured from five individuals per family at, or as close as possible to, 50% hatching for each family. Yolk-sac volume was calculated assuming the shape of an ellipse (Blaxter and Hempel, 1963):

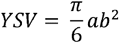

where a = length of the yolk sac (mm) and b = height of the yolk sac (mm).

### Statistical Analyses

Embryo survival was analyzed as a binomial response variable, while incubation period, length-at-hatch, and yolk-sac volume at hatching were analyzed as continuous response variables. Because embryos were raised independently, the replication unit in the statistical models is the individual embryo and the design was unbalanced due to different levels of embryo mortality. All non-proportional data were visually checked for approximate normality using histograms and Q-Q plots. A cubic transformation was applied to LAH and a cubic root transformation was applied to DPF, ADD, and YSV to normalize the distributions. Embryo survival was analyzed with binomial generalized linear mixed-effects models, and the transformed variables (i.e., DPF, ADD, LAH, and YSV) were analyzed with restricted maximum likelihood linear mixed-effects models with the *lme4* package v.1.1-26 (Bates et al., 2015). Population and incubation light treatment were included as fixed effects and female, male, female x male, and fertilization block as random effects. All traits and possible interactions were examined with backward, stepwise effect-selection and the maximal model for each trait selected using the *buildmer* package v.1.7.1 (Voeten, 2020). The significance for population, species, incubation temperature, interaction effects, and any random effects selected were determined using a likelihood ratio test between the maximal model and reduced models with the model effect of interest removed.

To enable population comparisons, the response to temperature for each trait was standardized to what we assumed was the optimal light treatment - the low light treatment (Table 1). For each trait and family, the within-family percent change from the optimal light intensity was calculated as:

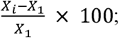

where x_1_ = mean trait value from low light treatment and x_i_ = the mean trait value from the light treatment of interest. The mean among-family percent change was calculated, and standard error was calculated as the among-family variation in percent change (Mari et al., 2021).

All analyses were performed in R version 4.0.4 (R Core Team, 2021).

## Results

### Spawning Adult and Egg Measurements

Lake Superior spawning adults ranged from 326-503 mm (total length mean (SD) = 412.5 (40.8) mm) and 298.9-970.0 g (fresh mass mean (SD) = 589.1 (171.4) g) and were larger in total length and fresh mass than Lake Ontario adults which ranged from 321-425 mm (mean (SD) = 372.5 (25.3) mm) and 280.5-795.8 g (mean (SD) = 496.6 (126.4) g). Egg diameter was larger in Lake Ontario (mean (SD) = 2.30 (0.08) mm) than Lake Superior (mean (SD) = 2.14 (0.12) mm).

### Developmental and Morphological Traits

Incubation period (both DPF and ADD) and YSV had significant interaction effects between population and light treatments (maximum *P* = 0.008; Table 3). The interaction effects precluded any interpretation of main effects for incubation period and YSV but did suggest contrasting norms of reaction between populations. Below we describe the interaction effects for incubation period and YSV, and the population main effects and light treatment pairwise comparisons for embryo survival and LAH. All random effects (i.e., female, male, and female x male) were significant (maximum *P* = 0.009) except female for embryo survival, male for embryo survival and YSV, and female x male for embryo survival and LAH (Table 3). All statistical model results can be found in Table 3.

**Table 3.**
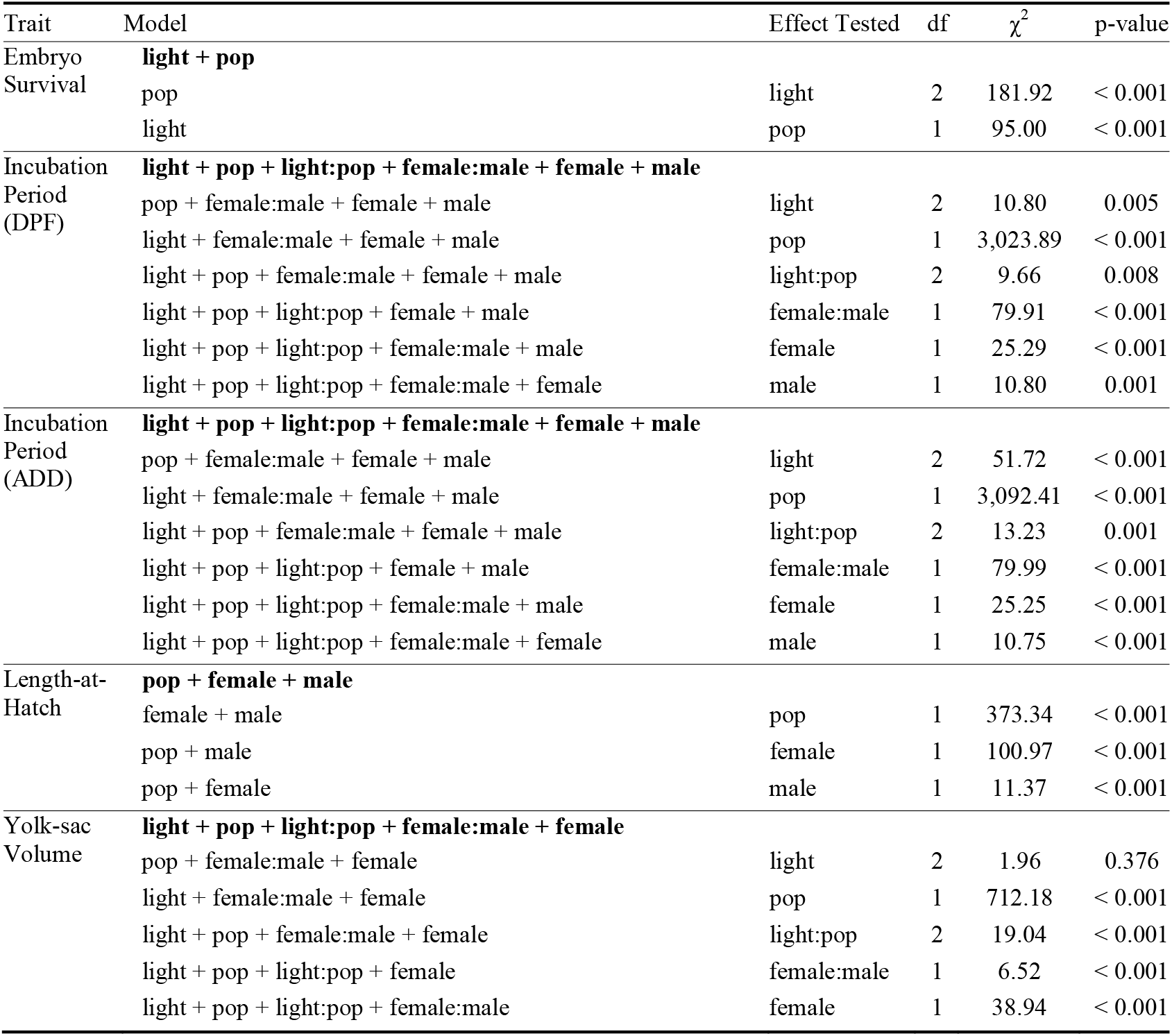
Likelihood ratio test output for each model selected for embryo survival (%), incubation period (number of days post-fertilization (DPF) and accumulated degree days (°C; ADD)), length-at-hatch (mm), and yolk-sac volume (mm^3^) from Lakes Superior and Ontario cisco (*Coregonus artedi*). pop indicates population. The full model that was selected is bolded for each trait.

### Embryo Survival

Embryo survival was highest for both populations at the medium light treatment, but lowest at the low light treatment for Lake Ontario and at the high light treatment for Lake Superior (Figure 3). Light and population main effects were significant. Only Lake Ontario pairwise light treatment comparisons with the low light treatment were significant (Low - Medium *P* < 0.001; Low - High *P* < 0.001). All pairwise light treatment comparisons for Lake Superior were not significant (minimum *P* = 0.089). Embryo survival was higher for Lake Ontario at the high (98.4%) and medium (99.6%) light treatments than for Lake Superior (85.3 and 89.3%, respectively) but there was no difference between populations at the low light treatment (0.9%; Figure 3).

**Figure 3.**
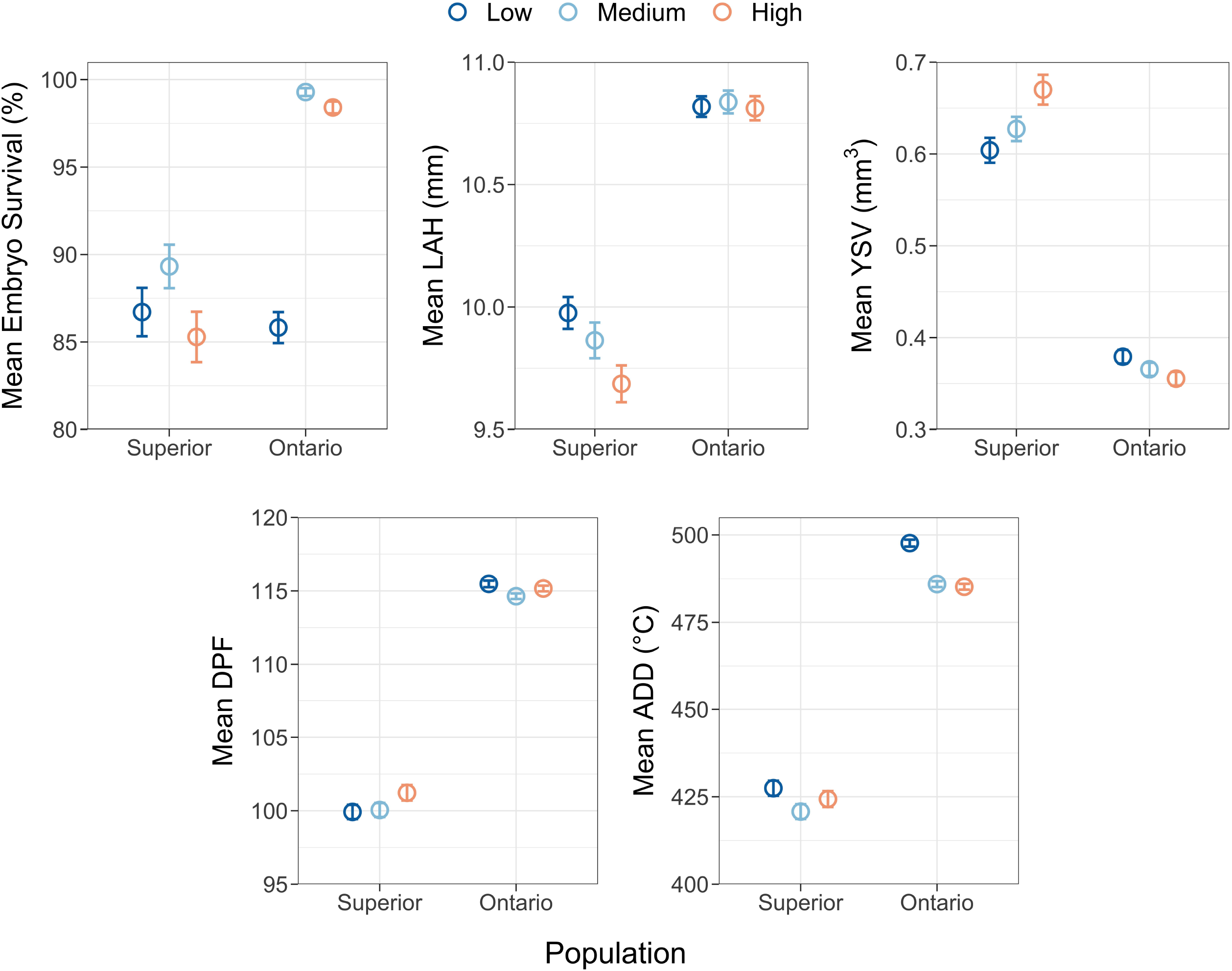
Mean embryo survival (%), length-at-hatch (mm; LAH), yolk-sac volume (mm^3^; YSV), and incubation period (number of days post-fertilization (DPF) and accumulated degree days (°C; ADD)) at each incubation light treatment from Lake Superior and Lake Ontario cisco (*Coregonus artedi*). Error bars indicate standard error.

### Incubation Period

The number of days between fertilization and hatching was highest for Lake Ontario at the low light treatment (115.47 days) and for Lake Superior at the high light treatment (101.22 days; Figure 3). Lake Ontario cisco had a decrease in DPF from the low light to the high light treatments (−0.7%), while Lake Superior had an increase from the low light to the high light treatments (1.9%; Figure 3). Incubation period (DPF) was longer for Lake Ontario than Lake Superior across all light treatments (mean (SD) difference = 13.9 (0.8) days).

The effect of light depended on population because the difference in ADD between populations was less pronounced at the high light treatment (difference = 60.8 ADD), while ADD was higher for Lake Ontario at the low and medium light treatments (497.7 and 485.9 ADD, respectively) than Lake Superior (427.5 and 420.8 ADD, respectively; Figure 3). Lake Ontario ADD had a negative response from the low to high light treatments (−2.5%), while ADD for Lake Superior did not change from the low to high light treatments (0.05%; Figure 3).

### Length-at-Hatch

Light was not a component returned in the stepwise-selected model for length-at-hatch, but the population main effect between Lake Ontario and Lake Superior was significant (*P* < 0.001; Table 3). Lake Ontario had a higher LAH than Lake Superior across all light treatments (Figure 4). Length-at-hatch decreased with increasing light by 3.2% in Lake Superior, but negligible differences in LAH were observed for Lake Ontario across light treatments (Figure 4).

**Figure 4.**
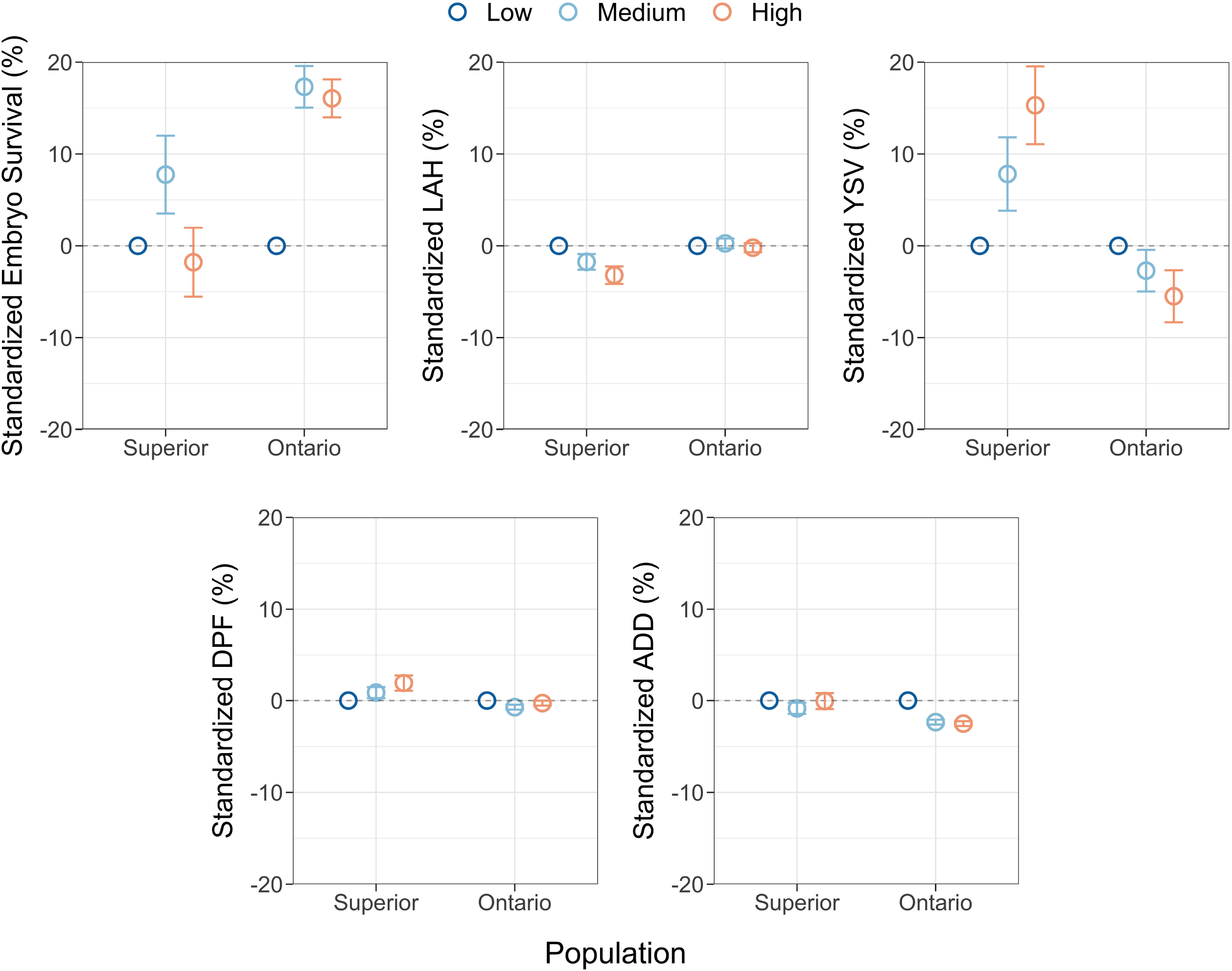
Mean among-family standardized responses (%) to assumed optimal light conditions (i.e., low) within each sampled population from Lake Superior and Lake Ontario cisco (*Coregonus artedi*) for embryo survival, length-at-hatch (LAH), yolk-sac volume (YSV), and incubation period (number of days post-fertilization (DPF) and accumulated degree days (ADD)). Error bars indicate among-family standard error.

### Yolk-sac Volume

Yolk-sac volume had a different response to light intensity between populations (Figure 4). The effect of light depended on population because the difference in YSV between populations was less pronounced at the low light treatment (difference = 0.22 mm^3^), while YSV was lower for Lake Ontario at the high and medium light treatments (0.35 and 0.37 mm^3^, respectively) than Lake Superior (0.67 and 0.63 mm^3^, respectively; Figure 4). YSV increased from the low to high light treatments (15.3%) in Lake Superior and decreased from the low to high light treatments (−5.5%) in Lake Ontario (Figure 4).

## Discussion

Developmental and morphological traits for Lake Superior and Lake Ontario cisco populations demonstrated similar and contrasting reaction norms to incubation light intensity. First, we found different responses to light intensity in embryo survival between populations. Second, increasing light intensity had minimal impact on incubation periods (DPF and ADD) for both populations. Lastly, LAH and YSV responded differently to varying light intensities between populations. Consequently, cisco from lakes Superior and Ontario are likely to have different responses to changes in ice coverage and subsequent light conditions.

Our hypothesis that embryo survival would be highest at the lowest light treatment was not supported. Embryo survival was highest for both populations at the medium light levels, suggesting that populations may be adapted to withstand some light exposure from high inter-annual variability in ice coverage. This result was contradictory to that observed in two Pacific salmonid species (chinook salmon *Oncorhynchus tshawytscha* and rainbow trout *Oncorhynchus mykiss*) for which embryo survival was highest at the lowest light exposures evaluated (0.04 μmol m^-2^ s^-1^; Eisler, 1961, 1958; Kwain, 1975). Lake Ontario cisco had a sharper decrease in survival than Lake Superior cisco at the low light treatment. The difference was surprising because average historical ice coverage over the Lake Ontario spawning location is higher than the Lake Superior spawning location, and thus low light conditions are more likely to occur for Lake Ontario cisco embryos. However, the Lake Ontario cisco spawning location is shallow (< 5 m) and could have high light intensity with little or no ice coverage. Higher variability in winter illuminance may have selected for the population of Lake Ontario cisco sampled to have greater flexibility to higher light conditions than deeper spawning cisco sampled from Lake Superior.

Our hypothesis that elevated light intensity would accelerate embryogenesis was not supported. The greatest difference in incubation periods was between populations, and was likely due to differences in embryo size, as larger embryos (i.e., Lake Ontario cisco) require more time to develop (Hodson and Blunt, 1986; Kamler, 2008). Previous studies of other salmonid species (European whitefish *Coregonus lavaretus*, chinook salmon, rainbow trout) found increasing light intensity decreased the length of incubation (Chernyaev, 2007; Eisler, 1958; Kwain, 1975).

In contrast to incubation period, LAH and YSV responded to the light treatment and matched our hypotheses, but responses differed between populations. Lake Ontario cisco exhibited minimal change in LAH as light increased, but YSV decreased, suggesting that light intensity increased the metabolic demand of embryos. In comparison, Lake Superior cisco showed a trade-off between LAH and YSV. A negative relationship between LAH and YSV is common in fish temperature incubation studies (Blaxter, 1991; Karjalainen et al., 2015; Stewart et al., 2021), but the relationship is usually accompanied by a change in incubation period as basal metabolic demand consumes yolk as a function of the length of incubation. We found that light influenced incubation periods similarly among light treatments; therefore, the trade-off between LAH and YSV in Lake Superior cisco suggests decreased yolk conversion efficiency to somatic tissue occurred as light intensity increased. This suggests future decreases in ice coverage and subsequent increases in embryonic light exposure, in the absence of adaptation, may result in smaller, less-robust larvae, which may in part explain the low survival of Lake Superior cisco and other coregonines to age-1 over the past 20 years (Lepak et al., 2017; Stockwell et al., 2009). The reasons underlying differences between cisco populations from Lakes Ontario and Superior remain unknown. However, the contrasting responses in LAH and YSV between populations suggests that embryogenesis for each population has different levels of developmental plasticity to light.

Embryo development is sensitive to environmental conditions, which can greatly influence life-history trajectories, performances, and reproductive success (Colby and Brooke, 1970; Karjalainen et al., 2016; Luczyński, 1991). We did not quantify developmental stages, except eye pigmentation, so specific life-stage developmental rates are unknown. Changes in the frequency of light (i.e., periodicity) can have adverse effects on fish embryos after yolk plug closure and first vertebrate trunk segment formation (Abdel-Rahim et al., 2019; Chernyaev, 2007, 1993; Ruchin, 2020). Fluctuating light cycles (e.g., 6:6h light:dark) and constant light (e.g., 24h light) accelerated the rate of embryonic development compared to ‘normal’ photoperiods (e.g., 12:12h light:dark; Chernyaev, 2007, 1993; John and Hasler, 1956; Ruchin, 2020). Photoperiod disruptions can inhibit the pineal organ and melatonin synthesis, which is critical to regulate and synchronize diurnal and seasonal biological rhythms (Delgado et al., 1987; Ekstrzöm and Meissl, 1997; Falcón et al., 2010; Roberts, 1978). The role photoperiod and the endocrine system plays in embryo development and phenology remains unknown for coregonines. Further studies that examine the impact of changing light intensities and photoperiods throughout incubations (e.g., decreased or no light during winter from ice coverage and increased light intensity and periodicity during spring ice-out) will help determine the fine-scale influence light and photoperiod may have on specific development stages (i.e., hatching), hormone regulation, and organ, tissue, and skeletal formation.

Sunlight intensity, albedo, and attenuation are strongly influenced by the angle of the sunlight, which is determined by season and latitude (Forsythe et al., 1995; Goldberg and Klein, 1977). Latitude and sun angle are negatively correlated and this negative relationship is strongest at the winter solstice in the northern hemisphere (Goldberg and Klein, 1977; Wielgolaski and Inouye, 2003). Lake Ontario is at a lower latitude and thus experiences a higher sun angle than Lake Superior, which results in a more intense and longer period of daylight. Our light treatments were calculated from light sensors deployed only in Lake Superior; thus, the experimental light intensity treatments for Lake Ontario cisco may not have captured an accurate light environment representation. Under-ice light data from more lakes, depths, and habitats would add to our understanding of cisco embryo light environments and improve the authenticity of experimental treatments. Additionally, comparing populations from high latitude lakes which experience decreased winter sunlight would provide an additional contrast for local adaptation and phenotypic plasticity across geographic regions.

Turbidity also contributes to light attenuation, and spring ice-out and river discharge can drastically increase the presence of suspended particulates and light absorption (Shao et al., 2019). Hydrological responses to climate change indicate earlier and protracted winter/spring runoff and higher runoff volume (Blahušiaková et al., 2020; Cutforth et al., 1999; Shen et al., 2018). Seasonal runoff, including snowmelt pulses, often drive high nutrient loads and primary productivity in temperate lakes (Isles et al., 2017; Rosenberg and Schroth, 2017). Runoff entering ice-covered lakes is expected to suspend near the ice surface, rather than mixing into the water column (Cortés et al., 2017; Yang et al., 2020), and therefore, have implications on when nutrients are used by primary producers and the onset of spring plankton blooms (Sommer et al., 2012). If the timing of spring plankton blooms changes as a result of earlier and protracted winter/spring nutrient loads, the potential mis-match between interacting species may produce bottom-up consequences (Rogers et al., 2020). Our results showed changing light intensities did not affect cisco hatch dates; thus, the ability of cisco to match optimal spring nursery conditions may be weakened if coregonines do not respond to changing ice conditions similarly to the plankton community (Cushing, 1990; Myers et al., 2015). The proximity of spawning and nursery grounds to shoreline and river outlets would likely impact the synchrony between coregonine larvae and planktonic prey.

Many fish species are iteroparous and, in some species, individuals repeatedly use the same spawning location (Marsden et al., 1995; Skjæraasen et al., 2011; Thorrold et al., 2001). The question of what constrains the choice of a spawning location cannot be separated from the question of what constrains early-life development and survival (Ciannelli et al., 2015; Iles and Sinclair, 1982; Petitgas et al., 2012; Sinclair and Iles, 1989). Embryo survival is largely determined by incubation habitat (e.g., water temperature, light exposure, oxygen availability, protection from predators), thus, selective pressure is focused on ‘correct’ and adaptive choices of spawning sites by the parents. The amount of spawning plasticity (e.g., spawning site selection, fidelity to spawning sites, spawning time) among populations could serve as an indicator for the level of evolutionary constraints for offspring (Ciannelli et al., 2015). For example, Atlantic herring (*Clupea harengus*) exhibit a wide range of reproductive strategies across diverse geographical locations, but have limited spawning site plasticity because embryo survival is dependent on substrate and vegetation (Petitgas et al., 2012). Coregonines are considered to be behaviorally and developmentally plastic and do not appear to be constrained by a spawning habitat type (Karjalainen et al., 2015; Muir et al., 2013; Paufve, 2019); however, our understanding of coregonine reproductive behavior and spawning-site selection is limited. The selection of deeper or shallower spawning locations would provide a gradient in environment conditions (e.g., light, temperature) depending on population-specific habitat requirements, and both suitable nearshore and offshore spawning habitats are historically likely to be present in each sampled lake (Goodyear, 1982; Paufve, 2019). Examining coregonine reproductive behavior and characterizing contemporary spawning habitat requirements is a logical and needed next step to build on our results.

The existence of varying trait responses between populations raises questions concerning causal mechanisms. Genomic studies can aid our understanding by determining what functional pathways could be up- or down-regulated due to light energy. Any potential changes in metabolic or catabolic genes from light will enhance trait analyses and allow further partitioning of the effects of light from other energy demanding environmental variables (e.g., temperature).

## Conclusion

Given the extensive degree of developmental plasticity in coregonines, propagation has been proposed as a practical way to reintroduce native species from lakes with extirpated or reduced population levels (Bronte et al., 2017; Zimmerman and Krueger, 2009). A key uncertainty to maximizing restoration efforts is whether managers should mimic natural environmental conditions to increase survival during propagation (Bronte et al., 2017). Our study highlights the potential role of winter light conditions, the influence of light intensity on cisco embryo development, and the impact changing ice regimes may have on cisco survival and recruitment in the wild. We did not identify a consistent directional reaction between and within the two sampled cisco populations to increasing light, and light is likely to have a differential effect on a number of physiological and biochemical processes. Large-scale, cross-lake propagation and reintroduction efforts are likely to be complicated by the capacity to match cisco phenotypes and optimal incubation conditions. Our results provide a step towards better understanding the recent high variability observed in coregonine recruitment and may help predict what the future of this species may look like under current climate trends.

## Acknowledgments

We thank the staff at the Wisconsin Department of Natural Resources Bayfield Fisheries Field Station, United States Geological Survey (USGS) Tunison Laboratory of Aquatic Science, and New York State Department of Environmental Conservation Cape Vincent Fisheries Station for conducting field collections of spawning adults. The staff at Apostle Islands National Lakeshore (U.S. National Park Service) conducted sensor deployment and retrieval. We also thank Rachel Taylor, Daniel Yule, and Caroline Rosinski for help with fertilizations and experiment maintenance. [NAME] provided the USGS solicited review that strengthened the manuscript, as did anonymous peer reviewers and Stockwell and Marsden lab members. This work was supported by the USGS [grant number G17AC00042]. Any use of trade, product, or firm names is for descriptive purposes only and does not imply endorsement by the U.S. Government.

